# Heterotrimeric PCNA Increases the Activity and Fidelity of Dbh, a Y-family Translesion DNA Polymerase Prone to Creating Single-Base Deletion Mutations

**DOI:** 10.1101/2020.06.08.140293

**Authors:** Yifeng Wu, William Jaremko, Ryan C. Wilson, Janice D. Pata

## Abstract

Dbh is a Y-family translesion DNA polymerase from *Sulfolobus acidocaldarius*, an archaeal species that grows in harsh environmental conditions. Biochemically, Dbh displays a distinctive mutational profile, creating single-base deletion mutations at extraordinarily high frequencies (up to 50%) in specific repeat sequences. In cells, however, Dbh does not appear to contribute significantly to spontaneous frameshifts in these same sequence contexts. This suggests that either the error-prone DNA synthesis activity of Dbh is reduced in vivo and/or Dbh is restricted from replicating these sequences. Here, we test the hypothesis that the propensity for Dbh to make single base deletion mutations is reduced through interaction with the *S. acidocaldarius* heterotrimeric sliding clamp processivity factor, PCNA-123. We first confirm that Dbh physically interacts with PCNA-123, with the interaction requiring both the PCNA-1 subunit and the C-terminal 10 amino acids of Dbh, which contain a predicted PCNA-interaction peptide (PIP) motif. This interaction stimulates the polymerase activity of Dbh, even on short, linear primer-template DNA by increasing the rate of nucleotide incorporation. This stimulation requires an intact PCNA-123 heterotrimer and a DNA duplex length of at least 18 basepairs, the minimal length predicted from structural data to bind to both the polymerase and the clamp. Finally, we find that PCNA-123 increases the fidelity of Dbh on a single-base deletion hotspot sequence 3-fold by promoting an increase in the rate of correct, but not incorrect, nucleotide addition and propose that PCNA-123 induces Dbh to adopt a more active conformation that is less prone to creating deletions during DNA synthesis.

**Highlights:**

- PCNA increases the fidelity of Dbh polymerase on a deletion-hotspot sequence.
- The interaction stimulates incorporation of the correct, but not incorrect, nucleotide.
- A minimal duplex length of 18 bp is required for PCNA to stimulate polymerase activity.
- Structural modeling suggests that PCNA induces a conformational change in Dbh.

## 1. Introduction

Y-family DNA polymerases are specialized DNA polymerases involved in translesion synthesis and DNA damage tolerance [1–3]. They are found in all three domains of life and are beneficial to cells because of their ability to bypass DNA damage that would cause the main replicative DNA polymerases to stall. When compared to the DNA polymerases that are responsible for the bulk of genome replication, the Y-family polymerases generally have a low fidelity of nucleotide incorporation. In cells, the activity of Y-family DNA polymerases is regulated both temporally and spatially so that lesions are bypassed without introducing excessive mutations into the genome.

The Y-family DNA polymerases interact with the sliding clamp processivity factor, which provides a way of targeting the polymerase to sites of stalled DNA replication [4,5]. In eukaryotes, the sliding clamp is called proliferating cell nuclear antigen (PCNA). In bacteria, the processivity factor is the β subunit of the Pol III replicative polymerase holoenzyme, called the β-clamp. Sliding clamps are all toroidal molecules that encircle the DNA duplex and have a pseudo six-fold symmetry [6–9]. The bacterial β-clamp is homodimeric, with each subunit having three similar domains [10,11]. Eukaryotic and archaeal PCNAs are trimeric, with each subunit having two similar domains. Although most of these are homotrimeric, a few eukaryotic and archaeal sliding clamps, including the human Rad9-Rad1-Hus1 (9-1-1) checkpoint complex [12,13] and the *Sulfolobus acidocaldarius* and *Saccharolobus solfataricus* (previously called *Sulfolobus sofataricus)* PCNAs [14,15], are heterotrimeric with three different subunits comprising the clamp.

The sliding clamp provides a docking platform for DNA replication and repair proteins and stabilizes their association with DNA. Binding is primarily mediated by a conserved motif in the partner protein, called the PCNA-interaction peptide (PIP) or β-clamp-binding motif, which binds to a hydrophobic pocket formed by the inter-domain connector loop (IDCL) of PCNA or β-clamp [16]. Each sliding clamp subunit has one binding site for other proteins, so that PCNA can bind up to three partner proteins simultaneously [15] and the β-clamp can bind up to two [17].

Binding to the sliding clamp is controlled in different ways in different domains of life. In eukaryotes, the Y-family polymerases bind to PCNA that has been mono-ubiquitinated [18], a modification that occurs as part of the DNA damage response. In bacteria, interaction of polymerases with β-clamp is largely governed by polymerase concentration and affinity of the PIP to the PCNA subunit [19]. Very little is known about how the archaea regulate the interactions between Y-family polymerases and PCNA, although the heterotrimeric complexes allow each subunit to bind preferentially to a different partner protein, providing another potential level of regulation.

Structurally, the Y-family polymerases all share a conserved right-hand configuration consisting of palm, thumb, and fingers domains, with an extra “little finger” or “polymerase-associated domain” (LF/PAD), and have no 3’-5’ exonuclease proofreading domain [20–23]. One significant structural feature of these enzymes is a spacious active site that enables the Y-family polymerases to accommodate various types of damaged DNA but which also reduces the fidelity of nucleotide incorporation [24]. Despite these structural similarities, each enzyme has a unique specificity for the types of lesions bypassed and fidelity of replication.

Two archaeal Y-family polymerases, DinB homolog (Dbh), from *S. acidocaldarius*, and DNA polymerase IV (Dpo4), from *S. solfataricus*, have been extensively characterized both structurally and biochemically. Even though these two polymerases share 54% sequence identity, they have distinctively different mutational signatures. In vitro, Dbh makes single-base deletion mutations at frequencies of up to 50% in specific deletion hotspot sequences, repeats of two or more identical pyrimidines flanked on the 5’ side by a guanosine (5’-GY_n_), but makes relatively few base-substitution mutations [25]. Dpo4, in contrast, has a much higher base-substitution mutation rate and a lower deletion rate, even though Dpo4 has a propensity to make deletion mutations in the same sequence context as Dbh [25–27].

Although Dbh and Dpo4 have been characterized extensively in vitro, much less is known about their functions in vivo. Inactivation of the *dbh* gene in *S. acidocaldarius* does not change the overall spontaneous mutation rate, but the mutation spectrum changes significantly, with fewer insertion-deletion mutations and more base-substitution mutations [28]. Additional studies indicate that Dbh is primarily responsible for the accurate bypass of 8-oxo-dG lesions, by correctly inserting dC opposite the lesion up to 90% of the time, but does not increase the frequency of single base deletion mutations in a 5’-GYn repetitive sequence that would be a deletion hotspot in vitro [29].

These findings suggest that Dbh is specifically recruited to sites of oxidative DNA damage and, because its access to replicating DNA is restricted, it does not introduce deletion mutations into the genome to the extent that the biochemical data would indicate. However, the intrinsic fidelity of Dbh may also increase in the cellular environment. Here we test the hypothesis that binding of PCNA to Dbh improves replication fidelity. We find that the physical interaction of Dbh with heterotrimeric PCNA-123 not only increases polymerase activity overall, but also decreases the single-base deletion rate.

## 2. Material and methods

### 2.1 Plasmid construction

DbhΔC10, a construct lacking the C-terminal ten amino acids of Dbh, was generated by PCR using a plasmid encoding full-length Dbh [23] as a template. The product was cloned into pET28a plasmid using In-Fusion PCR Cloning Kit (Clontech), with an N-terminal hexahistidine (His_6_) tag being added during the PCR amplification step. Genes for PCNA-1, PCNA-2, and PCNA-3, the three monomers of the heterotrimeric sliding clamp, were amplified by PCR from *S. acidocaldarius* genomic DNA (ATCC strain 33909) [30], and then individually cloned into pET28a plasmid, each with an N-terminal His_6_ tag added. PCNA-12 co-expression plasmids were constructed by inserting non-tagged PCNA-1 into multiple cloning site 1 and PCNA-2 (without and with a His_6_-tag) into site 2 of pET-DUET-1 plasmid (Novagen).

### 2.2 Protein expression and purification

Full-length Dbh was expressed in BLR(DE3)pLysS *E. coli* cells and purified by S-Sepharose chromatography as described previously [23,31]. DbhΔC10, PCNA-2, PCNA-3, and PCNA-12 expression plasmids were each transformed into Rosetta(DE3)pLysS *E. coli* cells. The transformed cells were grown in autoinduction media [32] at 37°C until the optical density at 600nm (OD600) reached 0.8. The temperature was then reduced to 20°C for overnight expression. Cell pellets were harvested the next day by centrifugation and resuspended in Buffer Ni-A (20 mM HEPES pH 7.5, 500 mM NaCl, 50 mM imidazole). For purification of the PCNA-123 complex, pellets from equal volumes of cell cultures expressing untagged PCNA-12 and His_6_-tagged PCNA-3 were mixed together prior to lysis. The PCNA-12 co-expression construct containing N-terminally His_6_-tagged PCNA-2 was used to purify these proteins in the absence of PCNA-3. Cells were lysed by sonication and then heated for 20 min at 75°C for DbhΔC10, 55°C for PCNA-123 and PCNA-12, or were not heated for PCNA-2 and PCNA-3. Subsequent steps in the purification were performed at 4°C.

Lysates were centrifuged for 1 hour at 20,000 × g, and the supernatant was loaded onto a HiTrap chelating HP column charged with nickel (GE Healthcare). Proteins were eluted from the column by a linear gradient (0 to 100%) of Buffer Ni-B (Buffer Ni-A with 500mM additional imidazole). Samples of eluted fractions were run on 15% SDS-PAGE gels. Fractions containing expressed protein were then pooled together and concentrated using either a 10 kDa, 30 kDa or 50 kDa Amicon Ultra-15 centrifugal filters (Millipore), depending on the size of the protein or protein complex. Elution buffer was also exchanged to protein storage buffer (20 mM HEPES pH7.5, 150 mM NaCl, 1 mM DTT) in the same step. Proteins were either kept at 4°C or were flash frozen with liquid nitrogen and stored at −80°C. Concentrations of the proteins were determined by UV absorbance at 280 nm using calculated extinction coefficients of 22,350 M^−1^cm^−1^ for Dbh and DbhΔC10, 17,880 M^−1^cm^−1^ for PCNA-2, 11,920 M^−1^cm^−1^ for PCNA-3, 38,280 M^−1^cm^−1^ for PCNA-12, and 50,200 M^−1^cm^−1^ for PCNA-123.

### 2.3 Size-exclusion chromatography

PCNA-2, PCNA-3, PCNA-12, or PCNA-123 were each mixed with Dbh or Dbh-ΔC10 in equimolar ratios, with each protein present at a final concentration of 200 *μ*M in a volume of 100 *μ*L. Proteins were then loaded on a Superdex 200 10/300 GL Tricorn gel filtration column (GE Healthcare), and eluted with 1.5 column volumes of gel filtration buffer (20 mM HEPES pH 7.5, 150 mM NaCl, 1 mM DTT). Fractions were collected and examined on a 4-20% pre-cast SDS-PAGE gel (Bio-Rad).

### 2.4 Primer-extension assays

Nucleotide incorporation assays were performed by rapidly mixing 50 *μ*L of reaction buffer (20 mM HEPES pH 7.5, 65 mM NaCl, 10 mM MgCl_2_, 1 mM DTT) containing polymerase (Dbh or Dbh-ΔC10), and primer-template DNA (with the primer 5’ end-labeled with 6-FAM fluorescein;

Integrated DNA Technologies) with an equal volume of reaction buffer containing one or more nucleotides (dATP, dCTP, dGTP, and/or dTTP) and incubated at room temperature for the times indicated in figure legends. When present, sliding clamp (PCNA-123 or PCNA-12) was mixed with the polymerase first, before other reaction components were added. After incubation, reactions were immediately quenched with the addition of an equal volume of reaction termination buffer (80% formamide, 100 mM EDTA, 4% SDS, 0.1% xylene cyanol dyes). Final concentrations of polymerase, clamp and nucleotide are indicated in the figure legends. When nucleotides were present at concentrations at or above 1 mM an equivalent concentration of MgCl_2_ was added to the nucleotide solution (in addition to the 10 mM MgCl_2_ in the reaction buffer), to keep the total concentration of free Mg^2+^ constant. At nucleotide concentrations 8 mM and higher, the activity of Dbh was inhibited (unpublished data, Y. Wu), making it difficult to obtain more accurate kinetic constants for experiment shown in Fig. 5 and Table 1. When used to prevent re-binding of Dbh to DNA, heparin was added to the nucleotide solution at a concentration of 1 mg/mL; control reactions in which heparin was pre-incubated with the protein and DNA at the same concentration showed no nucleotide incorporation demonstrating the effectiveness of the trap.

**Table 1.**
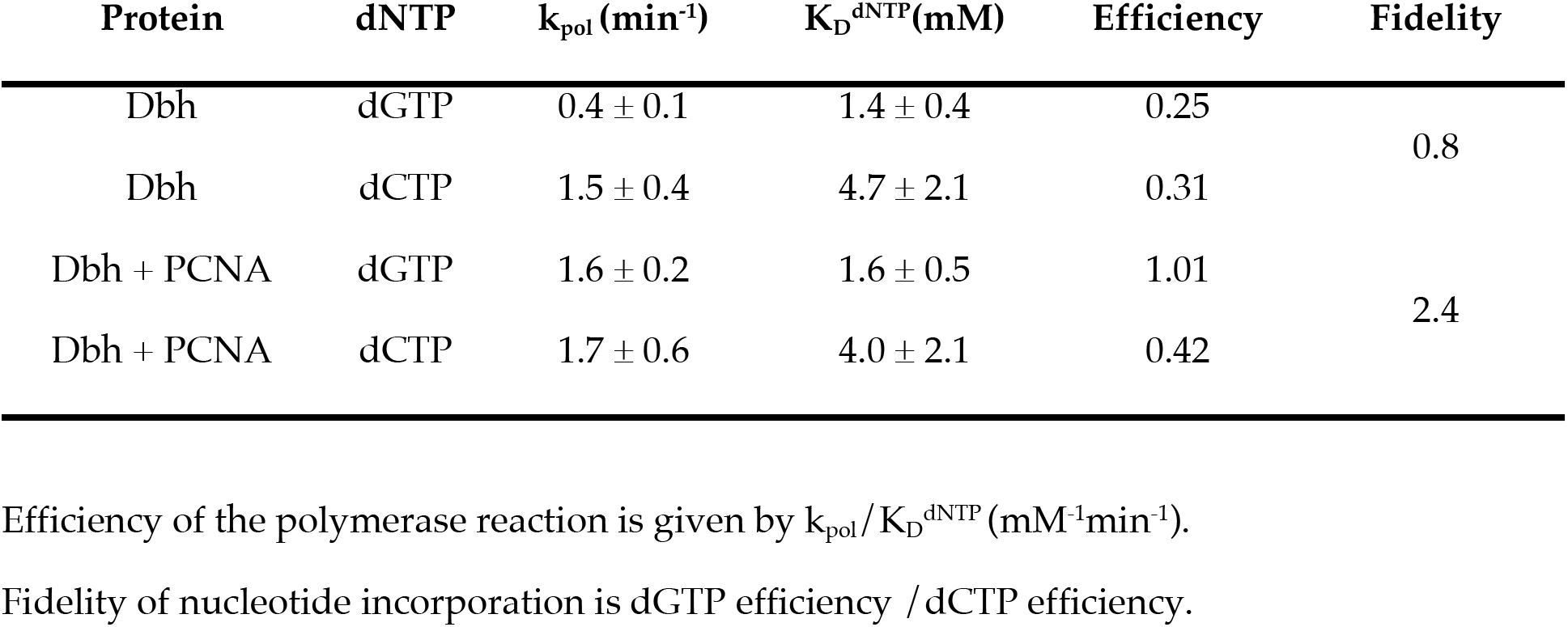
Nucleotide insertion kinetics for Dbh polymerase in the presence and absence of PCNA.

### 2.5 Gel electrophoresis and quantitation

Samples were incubated at 95°C for 5 min just prior to electrophoresis on 17.5% polyacrylamide (19:1), 7.5 M urea, 1× Tris-borate-EDTA (TBE) gels that were preheated and run at 50°C. The amounts of 6-FAM fluorescence in unextended and extended primer bands were quantitated using a Typhoon 9400 scanner and ImageQuant software (GE Healthcare). Single-nucleotide incorporation data were fit to the single exponential equation

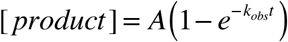

using GraphPad Prism software, where A is the amplitude, t is the time, and k_obs_ is the observed rate constant. The concentration of product formed was calculated from the fraction of primer extended and the concentration of primer-template DNA included in the reaction. The polymerization rate constant (k_pol_) and nucleotide dissociation constant (K_D_^dNTP^) were calculated by plotting k_obs_ against nucleotide concentration and fitting the data to the following hyperbolic equation using GraphPad Prism.

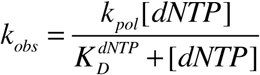

### 2.6 Structural modeling

PyMol version 2.3.5 was used to construct the models and to make the structural figures. A model of Dpo4 bound to dNTP, DNA and PCNA-123 was constructed as follows: first, the PCNA-1 subunit of the Sso PCNA-123 heterotrimer crystal structure (2NTI [33]) was superimposed on the PCNA-1 subunit in the crystal structure of Dpo4 bound to the Sso PCNA-12 heterodimer (3FDS [34]), creating a model of the complete clamp bound to the C-terminal amino acids of Dpo4 (residues 341-352); second, the C-alpha atom of Dpo4 residue 341 from a co-crystal structure of the polymerase with DNA and dNTP (3QZ7 [35]) was aligned with the same atom in the 3FDS structure and the coordinates of 3QZ7 were rotated around this point until the DNA was aligned towards the center of the clamp, positioning the enzyme so that it could both bind to the clamp and be actively replicating DNA; finally, the dNTP and DNA from the cryo-EM structure of human polymerase delta bound to PCNA (6S1M [36]) was aligned to the DNA of 3QZ7, via the alpha phosphates of the dNTP and the first 9 nucleotides of the primer strand (RMSD 1.16Å, 10 atoms), providing a 24-basepair (including the nascent basepair) B-form DNA duplex that extends through the clamp. This model (Fig. S1A) is similar to one previously published (see Fig. 5 in [34]), except that the DNA passes through the clamp at a more significant angle.

Based on the model of the actively replicating complex of Dpo4, dNTP, DNA, and PCNA-123, three alternative models with Dbh in place of Dpo4 were constructed. These models were based on aligning Dbh (3BQ1 [31]) with: (1) the LF / PAD of Dpo4 (residues 246-352; RMS, 0.50 Å, 82 atoms); (2) the palm domain Dpo4 (residues 1-20 and 78-170; RMSD of 0.39 Å, 90 atoms); and (3) the C-terminal tail of Dpo4 (residues 341-343). The latter modeling was possible to do because the crystal structures of Dbh shows two additional ordered residues at the C-terminal end of the LF/PAD than does the Dpo4 structure.

The first Dbh model (Fig. S1B) places residue 342 of Dbh 1.7 Å away from residue 341 of Dpo4 (the structurally equivalent residue), but the thumb of Dbh has a large steric overlap with the PCNA-2 subunit of the clamp and the DNA is more than 13 Å away from the polymerase active site (measured from the primer terminus to the active site residue D105). The second model of Dbh (Fig. S1C) aligns the DNA at the polymerase active site, but places residues 342 of Dbh and 341 of Dpo4 13.5 Å apart. The third model (Fig. S1D) aligns the overlapping three residues precisely and shows no steric overlap between the polymerase of Dbh and the clamp, but the LF/PAD overlaps the modeled DNA.

A final Dbh model (Fig. 6) was constructed by superimposing the coordinates of a chimera of Dbh (4NLG [37]) in which three amino in the linker region (243-KIP-245) were replaced with the structurally equivalent residues of Dpo4 (242-RKS-244) onto the structure of Dpo4 in the model described above (RMSD 0.956, 301 atoms). The position of Dbh in this model has no significant structural overlaps with either PCNA-123 or DNA.

A “supraholoenzyme” model (Fig. S2) of both Y- and B-family polymerases was constructed as follows: first, the co-crystal structure of Pfu DNA polymerase bound to a monomer of PCNA (PDB 3A2F, unpublished) was superimposed onto the PCNA-2 subunit of the Dbh model shown in Fig. 6, to model binding of an archaeal B-family DNA polymerase via the PIP to PCNA-123; then, the structure of *S. solfataricus* PolB1 (1S5J [38]) was superimposed onto Pfu DNA polymerase and Sso PolB1 was rotated to align the C-alpha atom of the last residue visible in the crystal structure (Ile 864) with the equivalent atom in Pfu (Lys 760). Next, the ternary complex structure of the B-family RB69 DNA polymerase (1IG9 [39]) was superimposed on the Sso PolB1 structure, to model binding of DNA, and finally, the structure of PolB1-DNA complex was manually adjusted so that the DNA pointed towards the center of the PCNA-123 heterotrimer while still maintaining the alignment of the terminus of PolB1 with the PIP of Pfu.

## 3. Results

Previously, the activity and processivity of Dbh had been reported to be stimulated by PCNA [40], however, only two of the PCNA subunits had been identified in *Sulfolobus* species at that time [41], so the significance of the findings has been unclear. The effect of the PCNA subunits from *S. solfataricus* on Dpo4 have been characterized previously [42], but because Dbh and Dpo4 have distinctly different biochemical activities, and because the PCNA 1, 2, and 3 subunits of *S. solfataricus* and *S. acidocaldarius* share 44.8, 48.8 and 54.5 % identity, respectively, we first characterized the interactions of Sac-PCNA-123 with each other and with Dbh, to determine if there were any significant differences between the proteins from the two organisms.

### 3.1 Dbh and PCNA physically interact through the C-terminal end of Dbh

We individually expressed and purified the *S. acidocaldarius* proteins Dbh, Dbh lacking the C-terminal 10 amino acids (DbhΔC10), PCNA-2, and PCNA-3 with multi-milligram yields of protein per liter of culture. PCNA-1 also expressed very well but was insoluble during purification. Co-expression of PCNA-1 with PCNA-2, however, yielded a soluble complex. A PCNA-123 complex was formed by mixing cells co-expressing PCNA −1 and −2 with cells expressing PCNA-3 prior to lysis.

Dbh and PCNA-123 interact and form a stable complex that can be isolated by size-exclusion chromatography (Fig. 1A). Equimolar concentrations of purified full-length Dbh and PCNA-123 were mixed and loaded on an analytical Superdex-200 gel filtration column. When the eluted fractions were analyzed, Dbh, PCNA-1, −2, and −3 were found together in one peak that eluted just after the 158 kDa marker. Since the molecular weight of Dbh is 40 kDa and of PCNA-123 is 90 kDa, and since each protein band is present in approximately equimolar amounts, we conclude that Dbh and PCNA-123 interact with each other and form a 1:1 stoichiometric complex. Some Dbh was also found in later-eluting fractions, consistent with some unbound polymerase. PCNA-1, −2, and 3 (but not Dbh), were also found in fractions eluting earlier, indicating that the sliding clamp subunits can form an even larger complex. Since the individual PCNA subunits were present in approximately equimolar amounts in the early-eluting peak, we suspect these fractions contain dimers of the PCNA-123 heterotrimer. A similar dimer was also found during purification of mammalian PCNA [43]. The absence of Dbh in these higher molecular weight fractions suggests that the putative dimeric form of PCNA-123 is in a conformation that blocks the site of Dbh interaction.

**Fig. 1.**
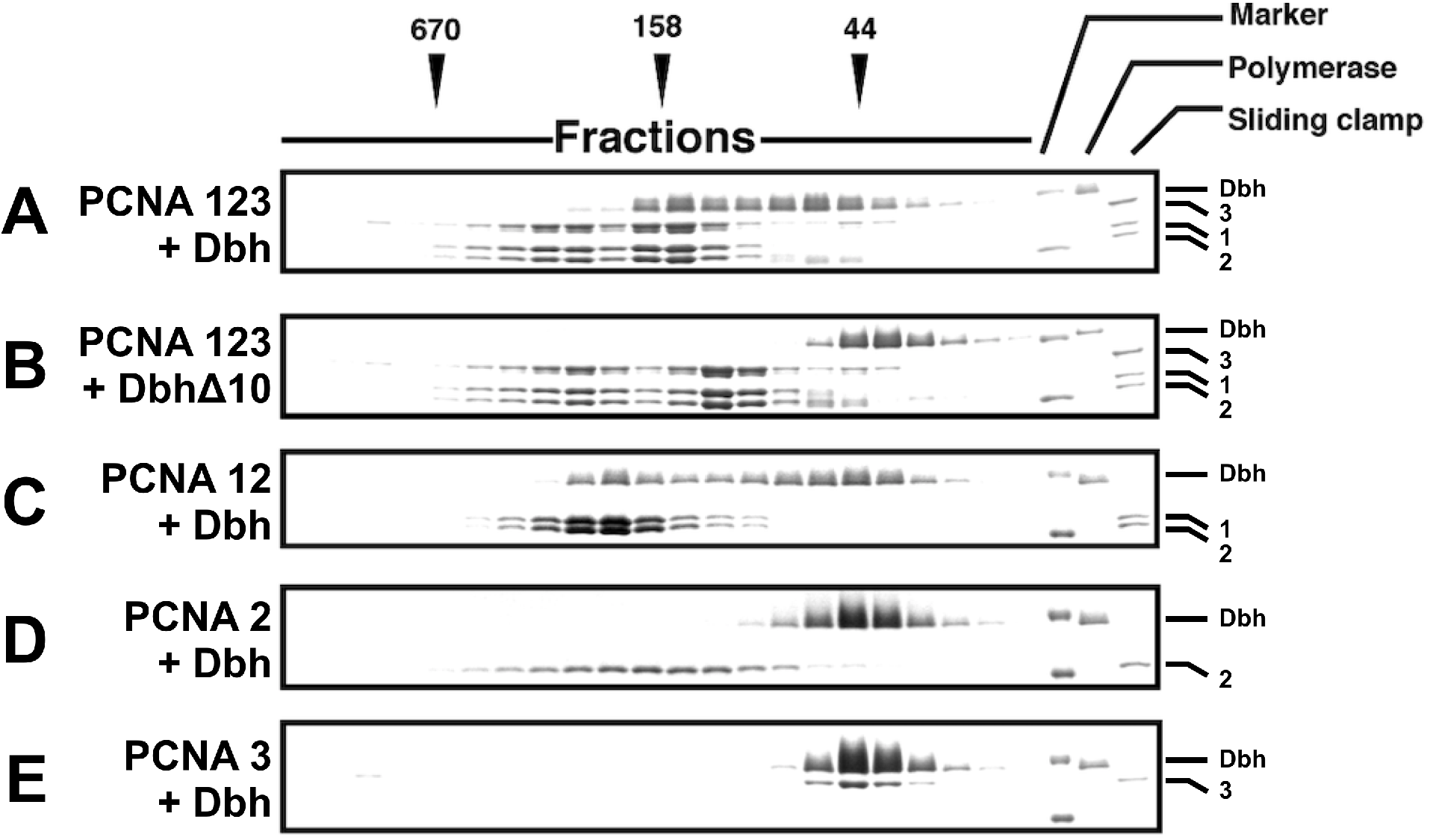
Dbh physically interacts with PCNA subunit 1. Fractions from size-exclusion chromatography of **(A)** full-length Dbh and PCNA-123; **(B)** DbhΔC10 and PCNA-123; **(C)** Dbh and PCNA-12; Dbh and PCNA-2; and **(E)** Dbh and PCNA-3. Elution positions of size-exclusion chromatography standards are indicated above the gels, with molecular weights given in kD. Molecular weights of the markers on the SDS-PAGE gel are 50 kD and 30 kD.

The C-terminal 10 amino acids of Dbh (345-<u>KTNLSDFF</u>DI-354) contain a putative PIP motif [44]. When these residues were deleted in DbhΔC10, the polymerase no longer co-eluted with PCNA, and instead eluted in fractions just after the 44 kDa molecular weight marker, consistent with the monomeric molecular weight of Dbh (Fig. 1B). The major peak containing PCNA-1, −2 and −3 (Fig. 1B) also eluted several fractions later than when the complex was formed with fulllength Dbh (Fig. 1A), indicating a smaller molecular weight of the PCNA complex, as would be expected in the absence of bound polymerase. As was observed in the case of full-length Dbh, some PCNA also eluted in a larger molecular weight peak (Fig. 1B). These results demonstrate that the C-terminal end of Dbh contains a functional PIP motif that is required for Dbh binding to PCNA.

### 3.2 Dbh interaction with the sliding clamp requires the PCNA-1 subunit

To determine which PCNA subunit(s) bind to Dbh, we mixed full-length Dbh with either the heterodimeric PCNA-12 complex or with the individual PCNA-2 and PCNA-3 subunits, then performed size-exclusion chromatography (Figs. 1C, 1D, 1E). Dbh co-elutes with both subunits of the PCNA-12 heterodimer (Fig. 1C). However, the complex eluted even before the 158 kDa size marker. The large size of this complex and the higher of intensity of the PCNA-1 and −2 protein bands compared to Dbh suggests that the complex may contain two copies of PCNA-12 and one copy of Dbh. A similar situation occurs for *S. tokodaii* PCNA, where two copies of PCNA-2 and two copies of PCNA-3 form a heterotetrameric ring structure [45]. When we mixed Dbh with PCNA-2, PCNA-2 eluted alone as a large complex, while Dbh eluted alone as a monomer (Fig. 1D). This indicates that Dbh interacts with PCNA-1 or to the interface formed between PCNA-1 and −2. It is also conceivable that PCNA-1 induces a conformational change that allows Dbh to bind to PCNA-2, but this would be a novel interaction mechanism. When Dbh and PCNA-3 were mixed together, both eluted together, but in a peak near the 44 kDa size marker, indicating that the two proteins do not interact (Fig. 1E). Our findings are consistent with a previous study reporting that Y-family DNA polymerase Dpo4 from *S. solfataricus* binds to PCNA-1 [46], the ortholog of PCNA-1 in *S. acidocaldarius* [47]. We conclude that the interaction of Dbh with PCNA-123 is dependent on PCNA-1, most likely through a direct interaction between the Dbh PIP and a canonical complementary site on the PCNA-1 subunit.

### 3.3 PCNA stimulates the DNA synthesis activity of Dbh, requiring the full PCNA-123 heterotrimer and the Dbh PIP motif

We determined how PCNA binding affects the overall DNA synthesis activity of Dbh using a linear primer-template DNA containing a 24 bp duplex (Fig. 2A). On its own, Dbh showed very low polymerase activity (Fig. 2B, left panel), as reported previously [23]. Even at the highest polymerase concentration tested (1 *μ*M), no full-length extension products were formed. When PCNA was included in the reaction at a concentration of 1 *μ*M, Dbh showed higher activity (Fig. 2B, right panel). At the highest concentration of Dbh, 1 *μ*M, some products were extended to the full length of the template DNA. We conclude from these assays that PCNA stimulates the overall DNA synthesis activity of Dbh.

**Fig. 2.**
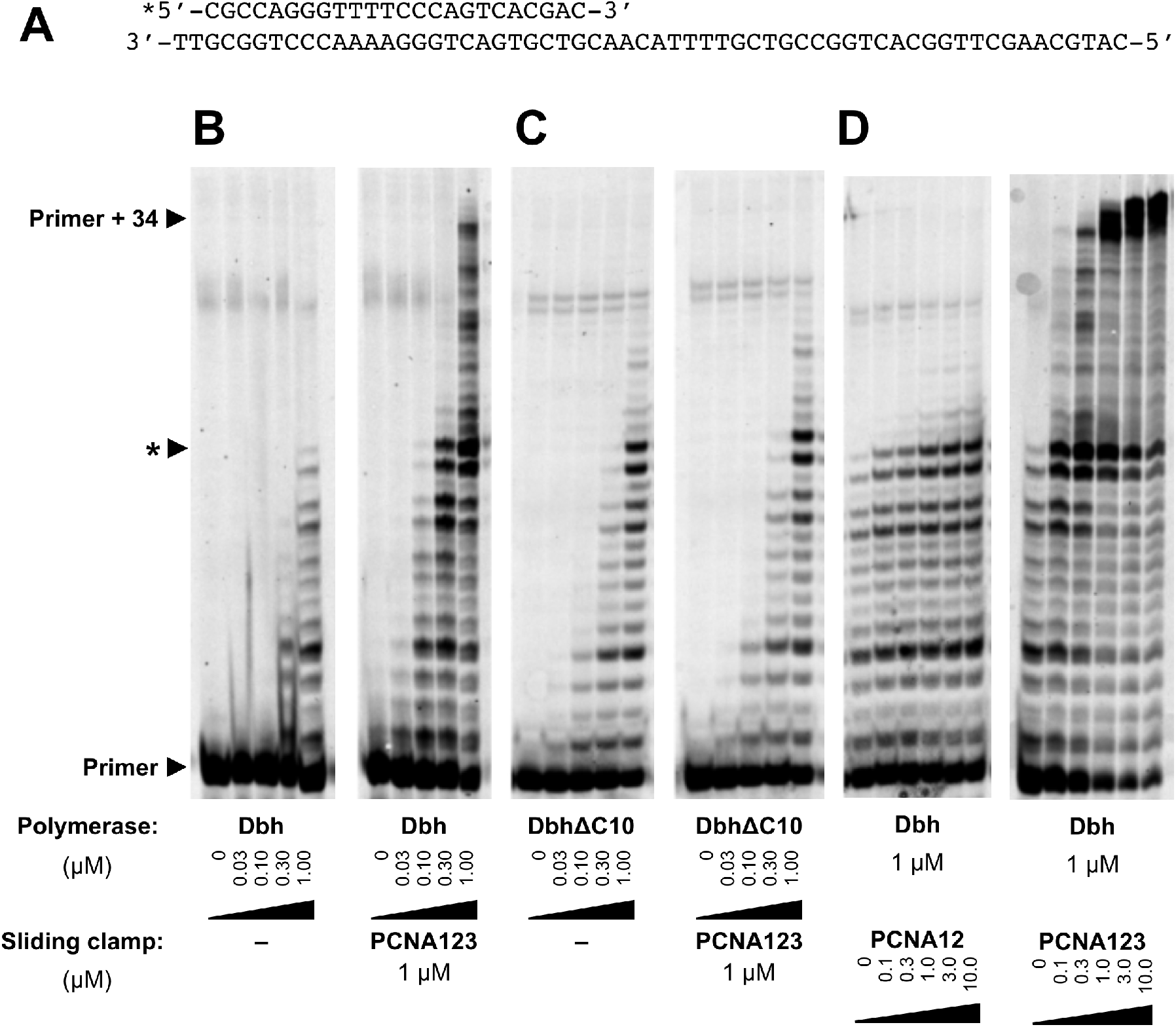
PCNA stimulates the overall DNA synthesis activity of Dbh. **(A)** Sequence of primertemplate DNA used at 1 *μ*M concentration in the assays. Multiple-nucleotide incorporation assays of increasing concentrations (0 *μ*M, 0.03 *μ*M, 0.1 *μ*M, 0.3 *μ*M, 1 *μ*M) of **(B)** Dbh and **(C)** DbhΔC10 without (left panels) and with (right panels) 1 *μ*M PCNA-123. **(D)** Increasing concentrations (0 *μ*M, 0.1 *μ*M, 0.3 *μ*M, 1 *μ*M, 3 *μ*M, 10 *μ*M) of PCNA-12 (left panel) or PCNA-123 (right panel) with 1 *μ*M Dbh. All reactions were incubated with 1 mM each of the four dNTPs for 20 min prior to electrophoresis on a denaturing polyacrylamide gel. A prominent site of stalling that is discussed in the text is indicated with an asterisk (*).

To test the importance of the PIP motif located at the C-terminus of Dbh, we repeated the same assay using the DbhΔC10 deletion mutant instead of full-length Dbh. Without PCNA, DbhΔC10 has a similar primer extension profile as full-length Dbh (Fig. 2C, left panel), but PCNA has no significant effect on the primer-extension activity of DbhΔC10 (Fig. 2C, right panel), with no full-length products being made in either case. This result, along with the result from the coelution assay, demonstrates that physical interaction between Dbh and PCNA through the PIPbox motif is required for enhancing the overall DNA polymerase activity of Dbh.

We also compared the effects of the complete PCNA-123 trimer and the PCNA-12 dimer complexes on Dbh. Using an increasing concentration gradient of each PCNA complex, we found that PCNA-12, shows very little stimulation of Dbh polymerase activity (Fig. 2D, left panel) compared to the intact PCNA-123 trimer (Fig. 2D, right panel). At a PCNA-123 concentration of 1 *μ*M, equal to the Dbh concentration in the assay, the majority of the primer was extended to the full length of the template. Only small increases in the amount of reaction products occur at higher PCNA-123 concentrations, consistent with a 1:1 stoichiometry between PCNA-123 and Dbh. In contrast, full-length primer-extension products are not observed even at the highest concentration of PCNA-12 (10 *μ*M) (Fig. 2D, left panel), comparable to the level of primer extension by DbhΔC10 in the presence of PCNA-123 (Fig. 2C, right panel). Thus, although Dbh binds to the PCNA-12 complex, a complete PCNA-123 heterotrimer is necessary to enhance the DNA polymerase activity of Dbh.

### 3.4 A duplex longer than 16 basepairs is required for PCNA to stimulate the activity of Dbh

Crystal structures have shown that Dbh contacts ~10 bp of primer-template DNA duplex [31] while β-clamp from *E. coli* encompasses ~10 bp of duplex DNA [10]. The assays reported in section 3.3 used substrate DNA with a 24 bp duplex region, which is sufficient for both the polymerase and clamp to bind to the DNA simultaneously. To determine if a minimal length of duplex DNA is required for PCNA to stimulate the polymerase activity of Dbh, we measured the rate of nucleotide incorporation on primer-template DNAs with duplex lengths of 16, 18, 20, and 22 bp (Fig. 3A). We incubated the DNA (40 nM) with a 100-fold excess of polymerase (4 *μ*M), to minimize the contribution of DNA binding to primer extension.

**Fig. 3.**
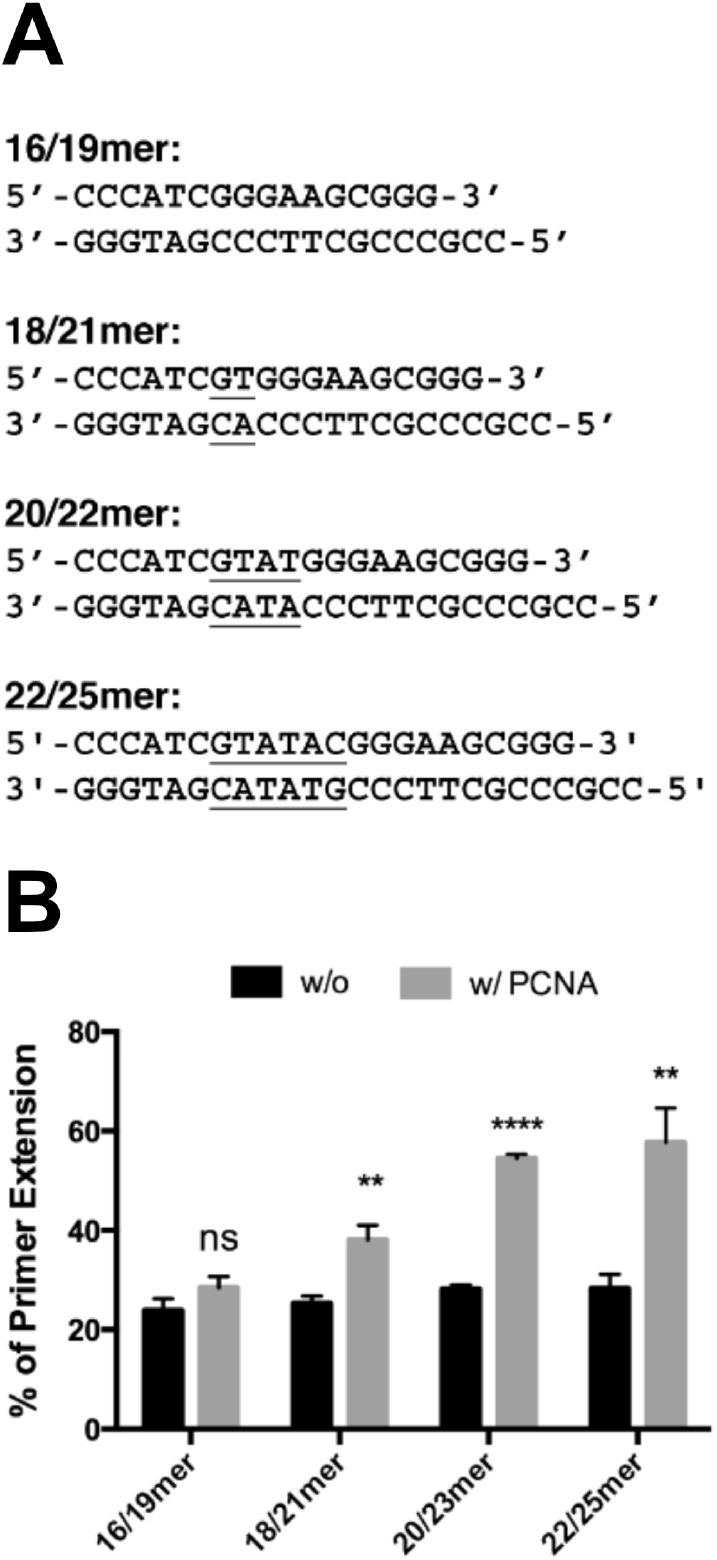
Stimulation of Dbh polymerase activity by PCNA requires a primer-template DNA duplex of at least 18 bp. **(A)** Sequences of primer-template DNAs with increasing duplex length. Nucleotides that vary between the sequences are underlined. **(B)** Primer-extension assays. For each substrate DNA, primer-extension reactions were performed under single-turnover conditions, using 4 *μ*M Dbh, 40 nM DNA, and 250 *μ*M dCTP, in the absence (black bars) or presence (gray bars) of 4 *μ*M PCNA-123. Reactions were terminated after 15 s and error bars indicate the standard deviation of measurements from three independent experiments. P-values are indicated for each DNA subsrate: P < 0.01, **; P < 0.0001, ****.

In the presence of PCNA-123, Dbh shows a duplex-length dependent increase in primer extension activity (Fig. 3B), with no significant stimulation on the 16-bp duplex length, and increases of 1.5-, 1.9- and 2.0-fold, respectively, on the 18-, 20- and 22-bp duplex lengths. In the absence of PCNA, Dbh extended all four substrates to an equal extent. We conclude that PCNA stimulates the activity of Dbh on substrates that are sufficiently long enough for the DNA to be encircled by the clamp.

### 3.5 PCNA stimulates the rate of nucleotide incorporation by Dbh

To fully assess the contributions of PCNA towards increasing the activity of Dbh, we followed the timecourse of DNA synthesis by Dbh in the absence or presence of PCNA-123 (Fig. 4), using the primer-template DNA shown in Fig. 2A that has a 24-bp primer-template duplex. Reaction products accumulate much more rapidly and reach full extension sooner in the presence of PCNA-123 than in its absence (Fig. 4, panel A vs. B). When the reactions were repeated in the presence of heparin, to prevent the polymerase from rebinding to DNA, no extension was observed (data not shown). A heparin trap is typically used to determine polymerase processivity, but under these conditions, Dbh is non-processive, with or without PCNA-123. This indicates that all of the products synthesized by Dbh are the result of multiple DNA binding events, with the rate of nucleotide incorporation (k_obs_) being slower than the rate of DNA dissociation (k_off_), even in the presence of PCNA-123. As a result, the probability of primer extension by polymerase for each encounter between the polymerase and DNA is less than one (k_obs_/k_off_ < 1).

**Fig. 4.**
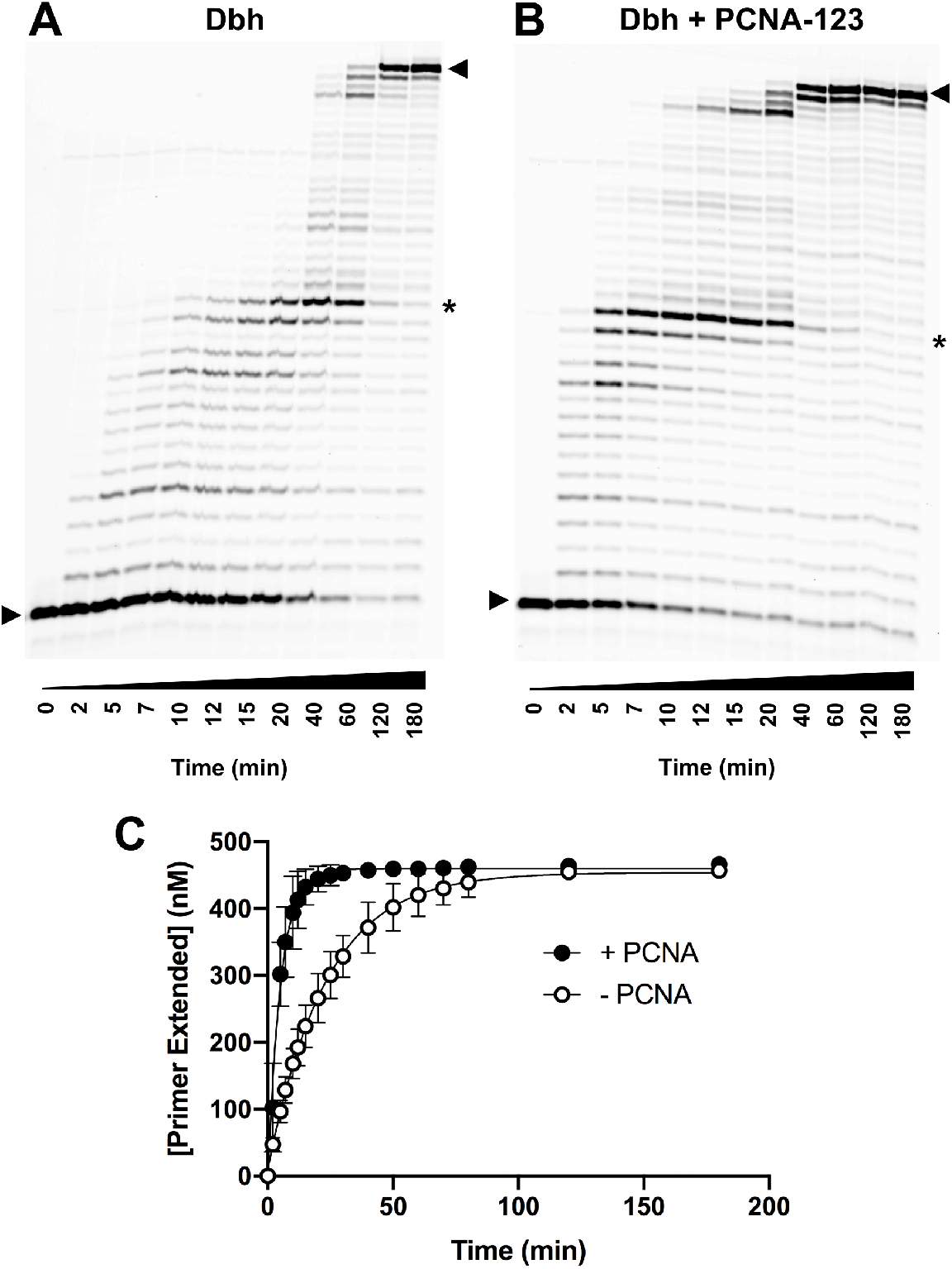
The presence of PCNA-123 increases the rate of nucleotide incorporation by Dbh. The primer-template DNA shown in Fig. 2A was extended by Dbh in the absence **(A)** or presence **(B)** of PCNA-123. Both annealed primer-template DNA and PCNA-123 were at a 500 nM concentration and incubated in the presence of 500 *μ*M each of the four dNTPs. The reaction was allowed to proceed for up to 180 minutes at room temperature, with aliquots being removed and quenched at the times indicated before being run on a 15% urea denaturing gel as shown. A prominent site of stalling that is discussed in the text is indicated with an asterisk (*). Triangles indicate the positions of starting primer and full extension product. The rate of incorporation of the single nucleotide dGTP is shown in **(C)**. Reaction conditions are identical to those in A and B, with the exception that a single nucleotide (dGTP) was provided. Once run on a 15% urea denaturing gel, the bands were quantitated with the results of product extension being plotted using Prism 8 to a single exponential. Error shown is standard deviation from two data points.

When Dbh is provided with just the first correct nucleotide, PCNA-123 stimulates the rate of incorporation by 5-fold, from 0.04/ min to 0.20/ min (Fig. 4C). These results indicate that PCNA-123 activates Dbh by a mechanism that is independent of its well know role in reducing the dissociation rate by topologically linking the polymerase to its template.

### 3.6 PCNA increases the fidelity of Dbh on a single-base deletion hotspot sequence

In the multiple nucleotide incorporation assays shown in Figures 2 and 4, there is a noticeable pause site where products accumulate after 14 nucleotides have been added. The template sequence in this region (5’-GCC-3’) corresponds to that of a predicted deletion hotspot (5’-GY_n_) [25]. To determine if PCNA might have any effect on the formation of single-base deletion mutations, we measured the activity of Dbh in single-nucleotide incorporation assays on the primer-template DNA shown in Fig. 5A, which contains the template sequence 5’-GCCC-3’ that we have characterized previously [37]. Observed rates of nucleotide incorporation (k_obs_) were determined for varying concentrations of the next correct nucleotide, dGTP, and the nucleotide that initiates a deletion, dCTP, in the absence and presence of PCNA-123 (Table 1 and Fig. 5B). The maximal rate of nucleotide incorporation (k_pol_) and the nucleotide dissociation constant (K_D_^dNTP^) were determined as described in the Materials section.

**Fig. 5.**
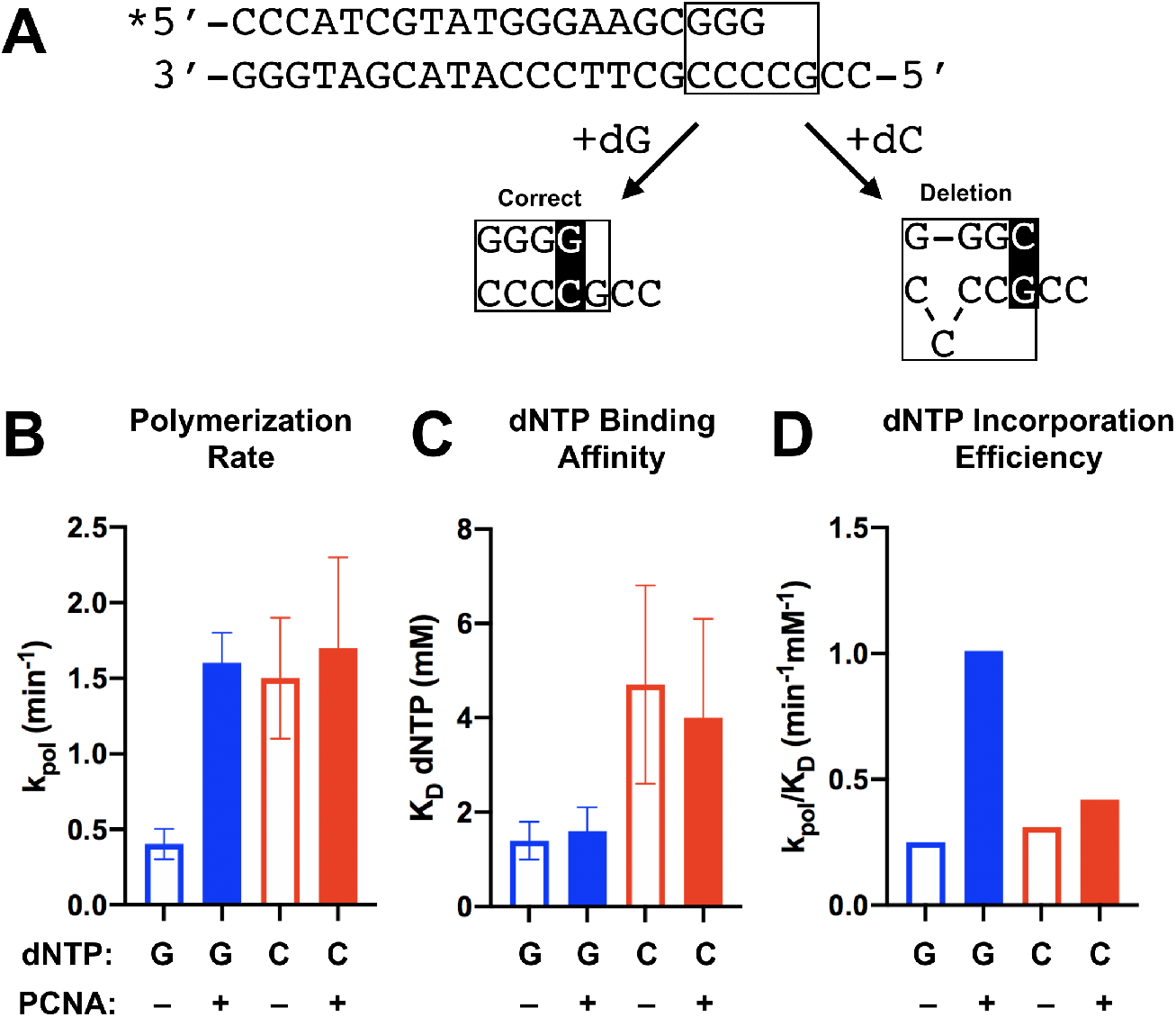
PCNA stimulates the incorporation of the next correct nucleotide by Dbh on a singlebase deletion hotspot sequence. **(A)** Sequence of the primer-template DNA used in the assay. The single-base deletion hotpot region is indicated with a box; the newly formed basepair is highlighted. Addition of dG yields the correct extension product while addition of dC initiates a deletion. Comparison of **(B)** polymerization rates (k_pol_), **(C)** nucleotide binding affinity (K_D_^dNTP^) and **(D)** nucleotide incorporation efficiencies (k_pol_/K_D_^dNTP^) for dGTP (blue) and dCTP (red) by Dbh in the absence (open bars) or presence (filled bars) of PCNA. Primer-extension assays used 4 *μ*M Dbh, 40 nM DNA and 0.1, 0.2, 0.5, 1.0, 1.5, 2.0 or 4.0 mM of either dGTP or dCTP and were performed in the absence or presence of 4 *μ*M PCNA-123. Observed rates of nucleotide addition (k_obs_) were calculated from progress curves of reactions that were incubated for varying times up to 60 min. Observed rates from two independent experiments were plotted against the nucleotide concentration and the data were fit to the hyperbolic equation given in the Materials and Methods to obtain the final kinetic and equilibrium constants shown in Table 1.

In the absence of PCNA, Dbh has a slight preference on this template for initiating a deletion rather than correctly extending the primer (Fig. 5B and Table 1). Dbh incorporates dCTP with a k_pol_ of 1. 5 min^−1^, ~4-fold faster than dGTP, which has a k_pol_ of 0.35 min^−1^. Binding of dCTP to Dbh is ~3-fold weaker than dGTP (K_D_^dNTP^ of 4.7 mM for dCTP vs. 1.4 mM for dGTP), resulting in preference of 1.2-fold (k_pol_/K_D_^dNTP^ of 0.31 for dCTP vs. 0.25 for dGTP) for Dbh initiating a deletion on this template sequence rather than incorporating the next correct nucleotide.

The fidelity of Dbh on this deletion-hotspot sequence, however, increases by 3-fold in the presence of an equimolar concentration of PCNA-123 (Fig. 5B and Table 1). The sliding clamp increases the rate of dGTP incorporation 4-fold (k_pol_ of 1.6 in the presence of PCNA vs. 0.4 min^−1^ in its absence) but does not substantially change the rate of dCTP incorporation (k_pol_ 1.5 min^−1^ vs. 1.7 min^−1^). PCNA had little effect on the binding affinity for either nucleotide. Thus, PCNA increases the replication fidelity of Dbh by increasing the rate of correct, but not incorrect, nucleotide incorporation.

## 4. Discussion

We have found that heterotrimeric *S. acidocaldarius* PCNA stimulates the activity of Dbh translesion polymerase, even on linear primer-template DNA where the sliding clamp is not topologically constrained. Enhanced polymerase activity requires the PIP-box motif of Dbh, a complete heterotrimeric sliding clamp, and a primer-template duplex long enough to be encircled by the clamp. Both Dbh and Dpo4 require the clamp-binding motif at the C-terminal end of the polymerase and both require the PCNA-1 subunit of the clamp, demonstrating the general conservation of the interaction. Dpo4 does not strictly require all three subunits of the clamp, as its polymerase activity can be stimulated by a construct in which the PCNA-1 and −3 subunits of the *S. solfataricus* clamp have been fused into one polypeptide, however it is not clear if these subunits would function in the same way if they were not fused [42]. Because the *S. acidocaldarius* PCNA-1 subunit is not soluble, we were not able to perform an equivalent experiment, but the heterodimeric PCNA-12 complex does not show any significant stimulation of Dbh activity.

A striking outcome of our studies is finding that PCNA enhances the fidelity of Dbh on a deletion hot-spot sequence as a consequence of the sliding clamp specifically increasing the incorporation rate of the next correct incoming nucleotide while having little effect on the incorporation rate of incorrect nucleotide. The fidelity of human pol η replicating an 8-oxo-G damaged template DNA base also increases in the presence of PCNA, but in that case, PCNA slows the rate of nucleotide incorporation overall and the increase in fidelity occurs because PCNA slows the rate of incorrect dATP incorporation more than it slows the rate of correct dCTP incorporation [48].

In contrast to the increased rate of correct nucleotide incorporation, PCNA did not significantly alter the affinity of Dbh for nucleotides (Table 1), although work on other Y-family polymerases indicates that this is possible. Increased nucleotide incorporation efficiency of *E. coli* pol IV in the presence of the β-clamp occurs as a result of an increased affinity of dNTP [49]. Similarly, eukaryotic PCNA also increased the nucleotide-binding affinities of yeast and human pol eta and of human pol kappa [50,51].

It is notable that PCNA enhances the activity and fidelity of Dbh for single nucleotide incorporation even under conditions where polymerase is in large excess over primer-template DNA. This suggests that increased DNA-binding affinity of the protein complex is not the basis for increased nucleotide incorporation efficiency. While PCNA may well stabilize binding of Dbh to DNA, we were unable to detect evidence of this in our experiments. We found that the rate of DNA dissociation must be faster than the rate of nucleotide incorporation both in the presence and absence of the clamp, indicating that Dbh is non-processive irrespective of the presence of the clamp. Our results are consistent with previous work showing that Dbh dissociates from DNA 10- to 100-fold more rapidly than rate of nucleotide incorporation [52]. In cells, where PCNA is unable to slide off DNA after being loaded, the processivity of Dbh is likely to be somewhat higher, but we would expect that the relatively slow rate of nucleotide incorporation, even at physiological temperatures [25], would still limit the amount of DNA synthesized by Dbh in vivo.

Several of our previous studies on the archaeal Y-family polymerases may help explain how PCNA increases both the activity of Dbh overall and the fidelity on deletion-hotspot sequences. First, Dpo4 from *S. solfataricus* has a higher nucleotide incorporation fidelity on the deletion hotspot sequence used here than does Dbh [31,35]. The fidelity of Dpo4 is higher because it incorporates the correct nucleotide more rapidly than Dbh, not because it incorporates the incorrect nucleotide more slowly. This is intriguingly similar to the observations reported here, where PCNA stimulates the rate of correct nucleotide incorporation by Dbh but does not significantly alter the rate of incorrect nucleotide incorporation.

Second, we have shown that the activities and nucleotide incorporation fidelities of Dbh and Dpo4 are largely determined by the inter-domain linker that connects the catalytic and LF /PAD domains [53]. Dbh is the less active of the two polymerases but has a higher nucleotide-incorporation fidelity in non-repetitive sequences than Dpo4. We found that these properties of Dbh and Dpo4 can be exchanged simply by exchanging the polypeptide linker that connects the catalytic and LF/PAD domains [53]. In fact, just three amino acids within the linker sequence are sufficient to determine these activities [37,53].

Third, the identity of the inter-domain linker is a major determinant of the overall conformation of the polymerase: the Dpo4 linker allows the LF/ PAD domain to contact the fingers of the catalytic domain while the Dbh linker restricts this movement [53]. Also, substitution of just three amino acids from the Dpo4 linker into Dbh allow the chimeric enzyme to adopt the standard conformation observed for Dpo4 [20]. Our crystal structures showed that primer-template DNA and dNTP are only positioned optimally for catalysis when the LF/PAD is in contact with the fingers.

Based on all these observations, we propose that the increased polymerase activity of Dbh occurs because PCNA induces a conformational change in Dbh that allows the DNA and incoming dNTP to be properly aligned at the polymerase active site for efficient catalysis. Induction of the conformational change would require PCNA-123 binding to the C-terminal PIP motif in Dbh and encircling the primer-template DNA duplex. We suggest that this conformational change would be inhibited by an extrahelical template DNA base, which occurs when single-base deletions are made [31,53], thus allowing PCNA to specifically favor the incorporation of the correct nucleotide and minimize the creation of deletions.

Despite many different attempts over the past three decades, no co-crystal structure has yet been determined of a sliding clamp bound both to a Y-family polymerase and to DNA. With regard to the work here, the most relevant available structures are of the LF / PAD domain of *E. coli* DinB (pol IV) bound to homodimeric β-clamp [54] and of full-length Dpo4 polymerase bound to PCNA subunits 1 and 2 from *S. solfataricus* [46], with neither structure having bound DNA. In both these structures, the polymerase (or polymerase fragment) is bound in an inactive conformation where it could not bind to primer template DNA as it passes through the central channel of the clamp.

We made use of the currently available structural information to create a model of Dbh actively replicating DNA in complex with PCNA-123 (Fig. 6). The key constraints used in building the model were that (1) the C-terminal tail of the enzyme that contains the PIP motif remained associated with the PCNA-1 subunit of the sliding clamp, as observed in the co-crystal structure of Dpo4 bound to PCNA-12 [34], but the enzyme could be rotated around the junction between the LF/PAD and the PIP motif, and (2) the DNA was both bound at the polymerase active site and passed through the central pore of the clamp. As described in the Material and Methods, we could not fulfill both of these requirements if Dbh remained in the same conformation observed in crystal structures [21,55]. If, however, the polymerase and LF / PAD domains were rotated relative to each other, as we have observed in the crystal structures of chimeric constructs of Dbh [37,53], we were readily able to construct a model that satisfied the constraints.

**Fig. 6.**
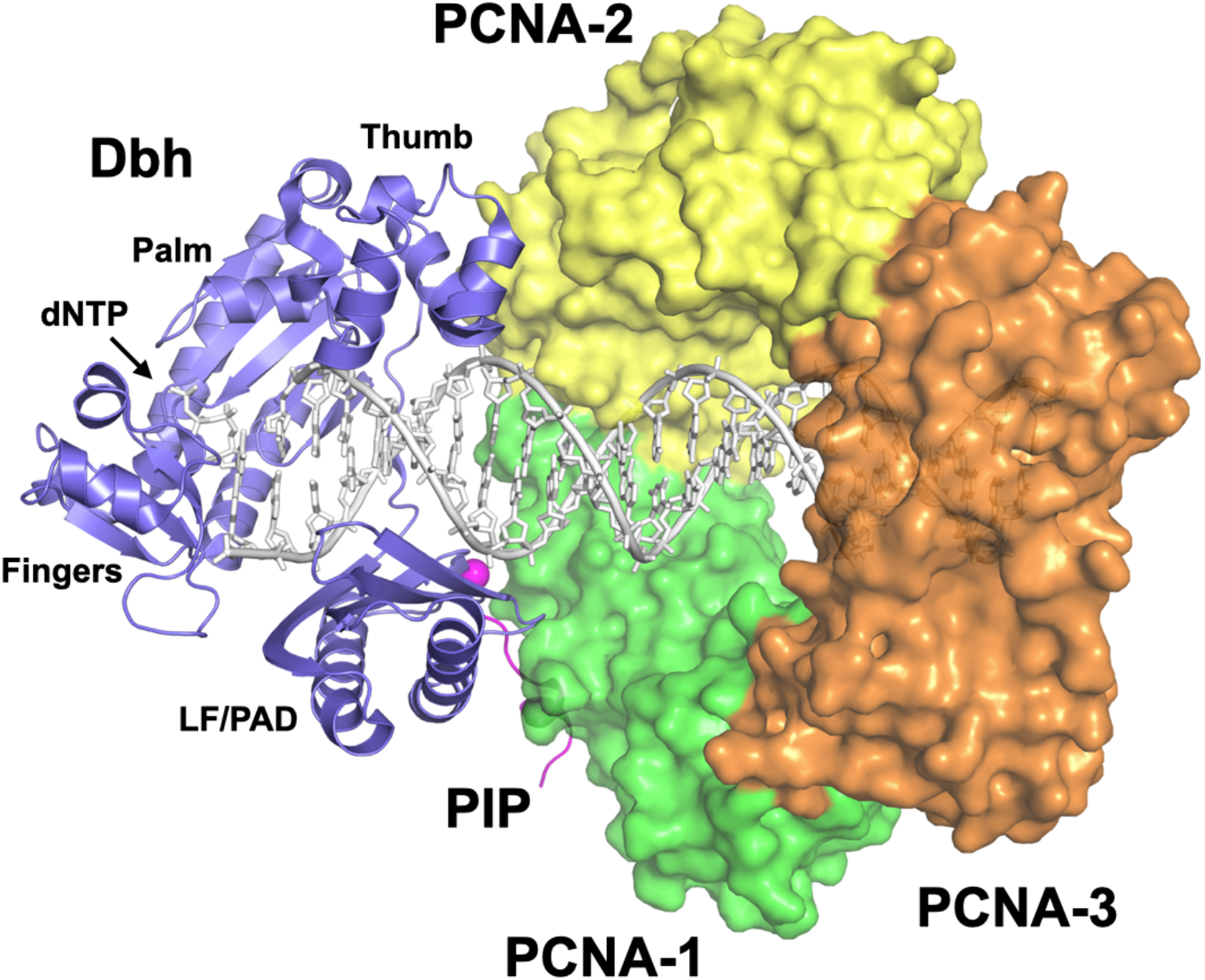
Model of an actively replicating Dbh-PCNA complex. Dbh must adopt the standard Dpo4 conformation to bind productively to the primer-template DNA and dNTP while avoiding steric clashes. The conformation of the Dbh polymerase (blue) is constrained by interactions of the PIP (magenta) with PCNA-1 (green), of the polymerase thumb with PCNA-2 (yellow), and for the DNA to pass through the clamp. See Materials and Methods for details. A 24-basepair DNA duplex (white) extends fully through the center of the sliding clamp. PCNA-3 is shown in orange and the C-alpha atom of Dbh residue 342 is shown as a magenta sphere.

An intriguing aspect of our model is that Dbh binding to PCNA-1 is fully compatible with a previous model of Sso PolB1 bound to PCNA-2 [56], forming the type of “supraholoenzyme” complex of B- and Y-family polymerases bound simultaneously to the PCNA-123 clamp (Fig. S2) that has previously been detected biochemically [57]. In the models of the individual polymerases, the DNA passes through the clamp at an angle and, in the supraholoenzyme model, the polymerase active sites face each other. Thus, polymerase switching could potentially occur simply by diffusion of the DNA between the active sites of the B- and Y-family polymerases, without any obstructions blocking the path.

In summary, the data presented here demonstrate that interaction of PCNA with Dbh increases the activity and fidelity of the polymerase. Structural modeling supports a mechanism by which the interactions between Dbh and PCNA facilitate a conformational change in the enzyme which promotes changes in polymerase activity that are consistent with the behavior of Dbh in cells.

## Acknowledgements

We thank Sean Fagan, Indrajit Lahiri and Purba Mukherjee for experimental advice and critical reading of the manuscript and Mona Gupta and Kim Hayden for assistance with protein purification. This work was supported by grant GM-080573 from the National Institutes of Health to J.D.P.

## Abbreviations

PCNA: proliferating cell nuclear antigen
TLS: translesion synthesis
ATP: adenosine triphosphate
dNTP: deoxynucleoside triphosphate
dGTP: deoxyguanosine triphosphate
dCTP: deoxycytidine triphosphate
PIP: PCNA interaction peptide
IDCL: inter-domain connecting loop1

---

**Fig. S1.**
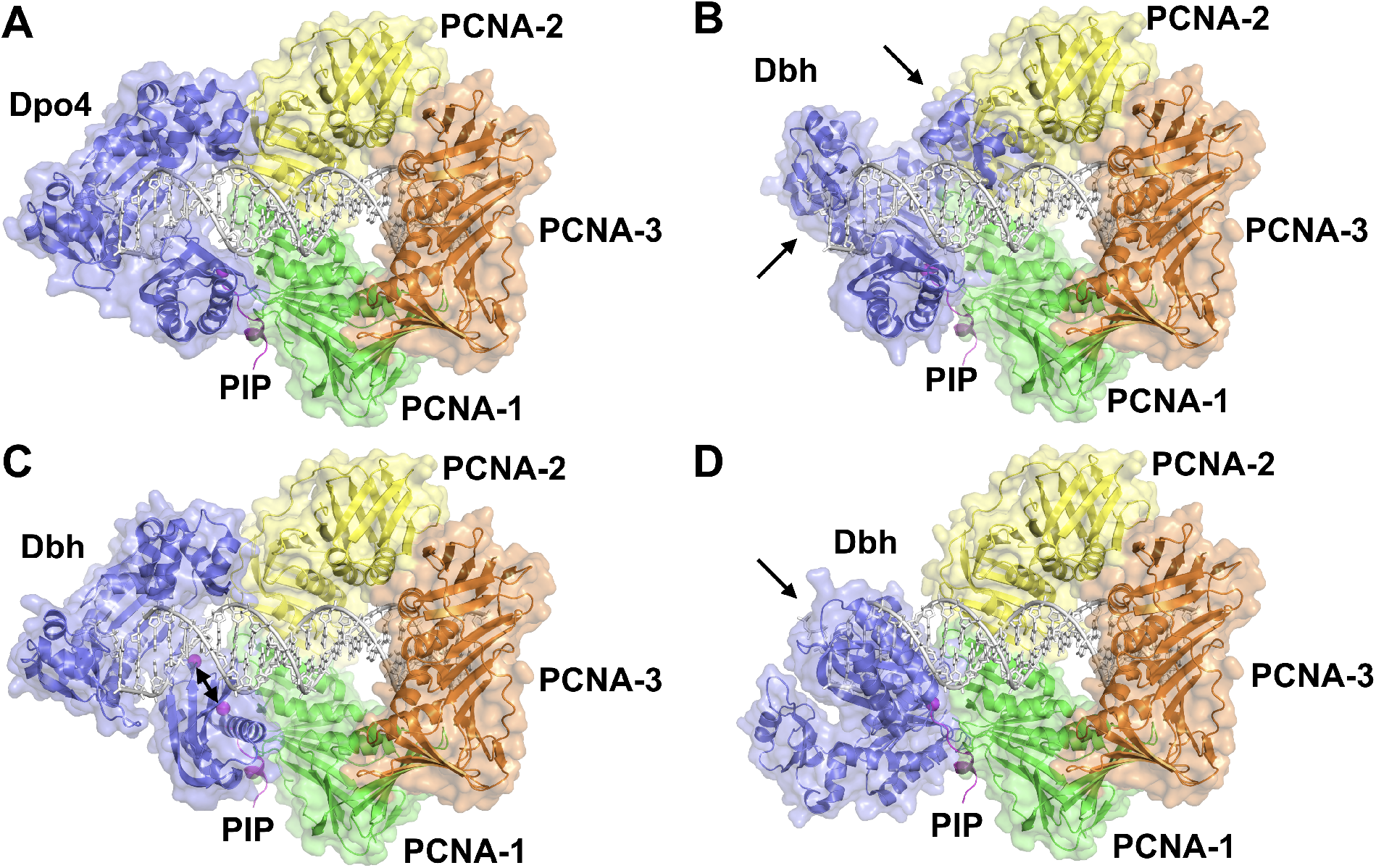
Models of Y-family DNA polymerases bound to dNTP, DNA and PCNA-123. The models show (A) an actively replicating complex of Dpo4, and models of Dbh based on superpositions with (B) the LF/PAD domain of Dpo4, (C) the polymerase domain of Dpo4, and (D) the PIP of Dpo4. In (B) and (D) single-headed arrows indicate steric clashes between Dbh and either PCNA-2 and/or the DNA. In (C) the double-headed arrow indicates the separation between Dbh residue 342 and the equivalent residue in the PIP. Details of the modeling are described in the Materials & Methods.

**Fig. S2.**
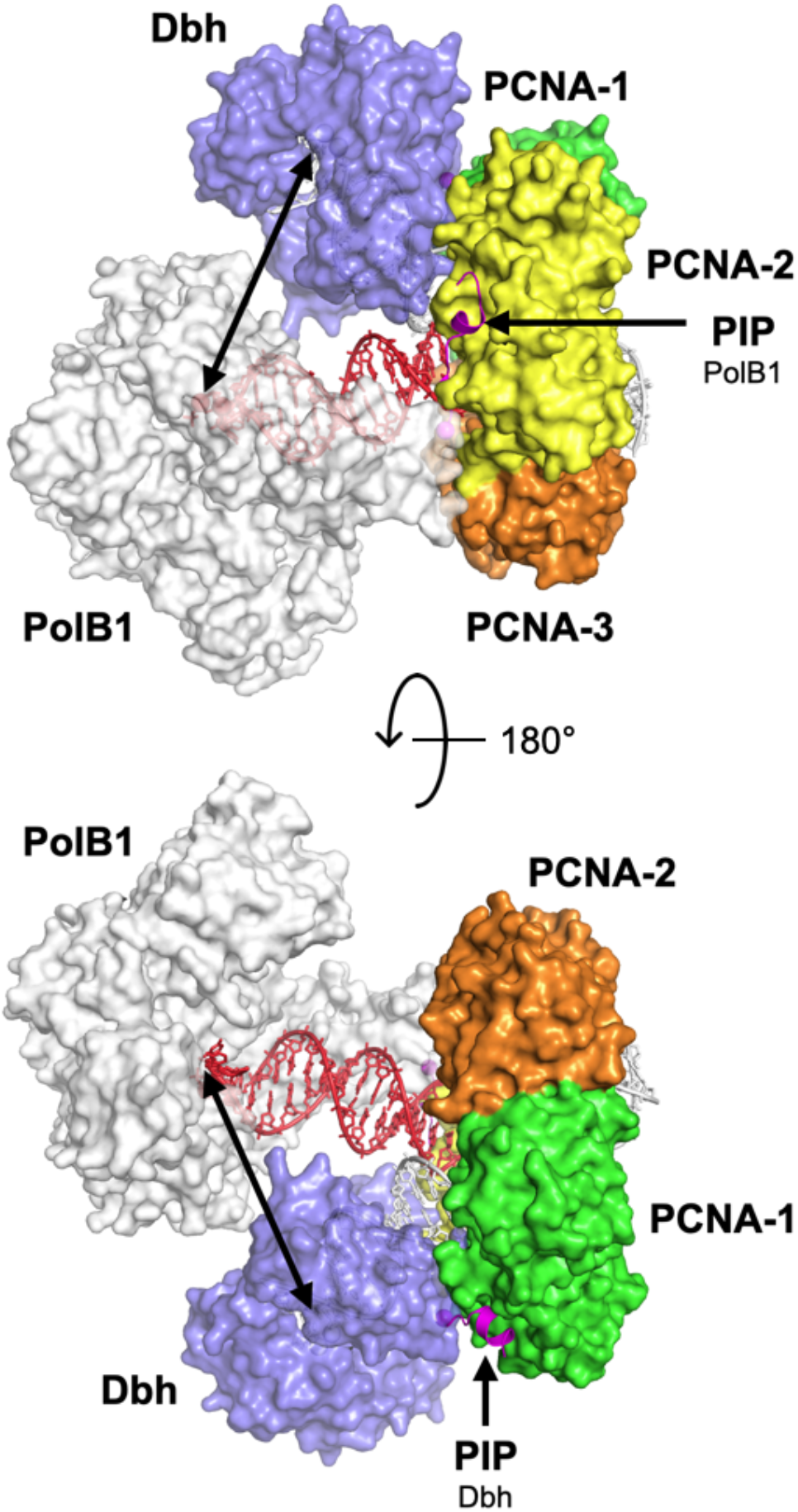
Model of “supraholoenzyme” complex of Y- and B- family DNA polymerases bound to PCNA-123. The model shows actively replicating complexes of Dbh and PolB1 in complex, with the PIP sequence of Dbh binding to PCNA-1 and the PIP sequence of PolB1 binding to PCNA-2. The path between the active sites of the two polymerases is indicated with a double-headed arrow. The top and bottom views are related by a 180° rotation around the axis indicated.

